# Androgen responsiveness to simulated territorial intrusions in *Allobates femoralis* males: evidence supporting the challenge hypothesis in a territorial frog

**DOI:** 10.1101/2020.11.15.383364

**Authors:** Camilo Rodríguez, Leonida Fusani, Gaëlle Raboisson, Walter Hödl, Eva Ringler, Virginie Canoine

**Affiliations:** Department of Behavioral and Cognitive Biology, University of Vienna, Vienna, Austria; Konrad Lorenz Institute of Ethology, University of Veterinary Medicine, Vienna, Austria; Messerli Research Institute, University of Veterinary Medicine Vienna, Medical University Vienna, University of Vienna, Vienna, Austria; Division of Behavioural Ecology, Institute of Ecology and Evolution, University of Bern, Bern, Switzerland

**Keywords:** Water-borne androgen, Challenge Hypothesis, poison frog, Androgens, Simulated territorial intrusion

## Abstract

Territorial behaviour has been widely described across many animal taxa, where the acquisition and defence of a territory are critical for the fitness of an individual. Extensive evidence suggests that androgens (e.g. testosterone) are involved in the modulation of territorial behaviour in male vertebrates. Short-term increase of androgen following a territorial encounter appears to favour the outcome of a challenge. The “Challenge Hypothesis” proposed by Wingfield and colleagues outlines the existence of a positive feedback relationship between androgen and social challenges (e.g. territorial intrusions) in male vertebrates. Here we tested the challenge hypothesis in the highly territorial poison frog, *Allobates femoralis*, in its natural habitat by exposing males to simulated territorial intrusions in form of acoustic playbacks. We quantified repeatedly androgen concentrations of individual males via a non-invasive water-borne sampling approach. Our results show that *A. femoralis* males exhibited a positive behavioural and androgenic response after being confronted to simulated territorial intrusions, providing support for the Challenge Hypothesis in a territorial frog.

## 1. Introduction

Territoriality is a widespread behaviour across many animal taxa and provides the territory holder with primary access to critical resources for individual fitness such as food, shelter, breeding sites and space for mating. In many species, only males engage in competitive interactions and contests with their conspecifics for the acquisition of territories (Davies, 1991). There is extensive evidence that androgens are involved in the modulation of typical territorial behaviours such as advertisement signalling and agonistic encounters in male vertebrates (Adkins-Regan, 2005). Testosterone is the main circulating androgen in most male vertebrates and, besides modulating the expression of primary sexual traits, its main function related to territoriality is to prepare males for social interactions, like male-male competition and agonistic encounters (Wingfield et al., 2006).

In most species with a seasonal breeding pattern, androgen levels undergo a seasonal fluctuation being higher during territory establishment and during the reproductive season. On the other hand, species with prolonged breeding and year-round territoriality present typical low androgen-baseline concentrations along the year but can facultatively rise during heightened male-male competition (i.e. territorial aggression towards intruders; Wingfield et al., 2006). The “Challenge Hypothesis” outlines brief increases of androgen levels in response to social challenges (i.e. territorial intrusions) in male vertebrates (Wingfield and Wada, 1989; Wingfield et al., 1990). This rapid increase in androgen levels promote aggressiveness, resource defence and mate guarding in a male-male competition context. Originally proposed for birds (Wingfield et al., 1990), the Challenge Hypothesis has been experimentally tested in fish, amphibians, non-avian reptiles and mammals (reviewed by Moore et al., 2019) by simulating a territorial intrusion (STI) test. These tests typically consist in presenting a stuffed or alive conspecific male decoy to a territorial male, combined /or solely with a conspecific acoustic stimulus, in order to quantify its aggressive response to this “intruder”. Results in different taxa were not consistent and had sometimes contrasting outcomes, prompting for a wider research across vertebrate taxa with a diverse suite of life-histories regulated by androgens (reviewerd by Moore et al., 2019). For instance, tropical amphibians provide ideal models for exploring the challenge hypothesis since they exhibit a multitude of strategies allegedly modulated by androgens such as parental care, sexual advertisement and/or territorial defence (Moore et al., 2005). So far, the few studies that investigated the challenge hypothesis in amphibians yielded contrasting results. For instance, in males of the Smith frog (*Hypsiboas faber*) testosterone levels did not increase after challenging males with STIs (de Assis et al., 2012). Otherwise, in the túngara frog (*Engystomops pustulosus*), water-borne testosterone concentration increased after confronting males to a combined acoustic and chemical (excretions of calling males) stimulus simulating a male competitor (Still et al., 2019).

Almost all male Neotropical poison frogs (Dendrobatidae) have been found to exhibit pronounced territoriality, showing aggressive territorial defence towards conspecifics (Pröhl, 2005). To date, it is not clear whether male territoriality in poison frogs is modulated by androgens. Part of this uncertainty is due to limitations in the collection of tissue for hormone measurement in small anurans. Classical hormone measurement methods are based on blood samples (Narayan, 2013) because hormones are systemic signals primarily released into the blood stream from the endocrine system. However, blood sampling may be difficult in small animals because of the amount of plasma needed for hormone quantification. Additionally, blood sampling usually requires prolonged handling of animals and invasive sampling procedures, which can influence the hormonal response and therefore affect the interpretation of the results obtained in experiments (Fusani et al., 2005; Hau and Goymann, 2015; Romero and Reed, 2005). Recently, water-borne sampling has been validated for measuring multiple steroid hormones from a single water sample (Baugh and Gray-Gaillard, 2020; Baugh et al., 2018; Gabor et al., 2016, 2013). By reflecting plasma steroid concentrations through its metabolic products, water-borne sampling has become an advantageous and non-invasive technique that minimizes the stress to the animals and allows the researcher to repeatedly measure hormone levels in the same individuals.

In this study, we tested the effect of territorial challenges on the behavioural and androgenic response of the Brilliant-thighed poison frog, *Allobates femoralis*. This species has become a model system for the study of acoustic communication (Amézquita et al., 2006, 2005; Gasser et al., 2009; Rodríguez and Hödl, 2020), spatial navigation (Pašukonis et al., 2016, 2014a, 2014b, 2013) reproductive (Ringler et al., 2018; Stückler et al., 2019; Ursprung et al., 2011) and social behaviour (Narins et al., 2003; Ringler et al., 2017; Rodríguez et al., 2020; Tumulty et al., 2018) in poison frogs. Males vocally advertise and actively defend their territories to conspecific males (Hödl, 1983; Narins et al., 2003; Ringler et al., 2011; Roithmair, 1992). No previous research has explored the proximate mechanisms underlying territorial behaviour and social interactions in *A. femoralis*, but it is likely that androgens (i.e. testosterone) modulate its calling and territorial behaviour as this appears to be an evolutionary conserved mechanism across vertebrate taxa (Simon and Lu, 2006; Wingfield et al., 2006).

In order to investigate whether a territorial intrusion induces an increase in androgens levels of *Allobates femoralis* males (as expected following the Challenge Hypothesis), we challenged territorial males performing STI-tests by presenting a playback stimulus. This method has been successfully used to induce a territorial response (i.e. positive phonotaxis) in *A. femoralis* males (Narins et al., 2006; Ursprung et al., 2009). Additionally, we measured males’ pre-and post-challenge androgen concentrations from water samples. Prior to analysis, we carried out a series of laboratory tests to investigate if hormonal concentrations in the holding water relate to those in the blood. Since there might be effects of time of the day and spontaneous behaviours on androgens concentration (Wada, 1986), we measured calling, locomotor, courtship and foraging activity across the day in addition to water-borne androgen levels prior to the territorial challenge. We further examined whether the intensity of the behavioural response was coupled with the androgen response to STIs.

## 2. Methods

### 2.1. Study system

The brilliant-thighed poison frog, *Allobates femoralis*, is a diurnal and terrestrial species belonging to the family Dendrobatidae (“AmphibiaWeb,” 2020; Boulenger, 1883). Males exhibit strong territoriality within the prolonged breeding season, which usually begins with the onset of the rainy season (Kaefer et al., 2012; Montanarin et al., 2011). During territorial interactions or courtship displays, males produce acoustic signals from elevated structures on the forest ground. Courtship behaviour is accompanied by a locomotor display called “courtship march”, which usually starts in the afternoon and ends on the next morning (Hödl, 1983; Ringler et al., 2013; Roithmair, 1992; Stückler et al., 2019). Males’ territorial displays consist, first, in antiphonal calling to warn neighbouring males of the ownership of a territory, and second, in a phonotactic or agonistic response towards intruder males (Hödl, 1983; Narins et al., 2003; Roithmair). Vocal behaviour of *A. femoralis* males is more frequent in the afternoon than in the morning (Roithmair, 1992).

### 2.2. Sample collection and hormones extraction

#### 2.2.1 Water-borne hormone sampling and solid-phase extraction (SPE)

For water-borne hormone sampling and extraction we followed published methodology (see below). It is noteworthy that by measuring anti-androgen immunoreactive substances in water samples we cannot exclude some of androgenic conjugate forms (Baugh and Gray-Gaillard, 2020). Therefore, along this manuscript we refer to “water-borne androgen” by actually referring to androgens and metabolic products in the holding water, as it is mentioned in similar publications (Scott et al., 2008). Every water bath consisted of a glass container (14cmx9cmx5cm) filled with 40 mL of distilled water. Frogs were placed in the water bath immediately after capture and removed after 60 min, and then released at the original location. Androgens were extracted by collecting each water sample with 20 mL sterile syringes coupled to an individual C18 cartridge (SPE, Sep-Pak C18 Plus, 360 mg Sorbent, 55 - 105 μm particle size, #WAT020515, Waters corp., Milford, MA) with a flow rate of ca. 10 mL/min. Later on, cartridges were eluted with 4 mL of 96% EtOH into 8 mL borosilicate vials and stored at 4°C until further hormonal analysis in laboratory. Samples were dried down with N2 at 37°C, resuspended with 250 uL of the assay buffer (provided in the ELISA kit, see below) and incubated overnight at 4°C.

In order to calculate recovery efficiency of testosterone with the SPE technique, we spiked two pools with 2 different testosterone concentrations, using standards of the ELISA kit (see below). Samples were extracted and processed as described above and stored at 4°C until proceeding with the assay. Water samples “without frog” were also processed as blank controls to evaluate any possible contamination of the holding water. To assess water-borne androgen release rate, we used thirteen adult *A. femoralis* (Body-size mean ± SD: males=2.74 ± 0.03 cm, N=7; females = 2.79 ± 0.02 cm, N=6) in January 2018 from a laboratory population kept at the animal care facilities at the University of Vienna. Briefly, we manually placed each frog in consecutive water baths of sampling periods of 15, 30 and 60 min. All samples were collected between 08:00 and 09:00 A.M., then extracted and processed as described above and, stored at 4°C until the assay. All frogs were fed at libitum with wingless fruit flies every second day.

#### 2.2.2. Parallelism between hormone concentration in blood and holding water

In order to know whether water-borne androgen reflected actual levels of circulating testosterone at the time of sampling, we collected eighteen free-living adult *A. femoralis* males (Body-size mean ± SD = 2.8 ± 0.1 cm) in April 2019, from a population in the vicinity of Roura, French Guiana (4°43’ N – 52°18’ W). Frogs were attracted using playbacks, captured using plastic bags and transferred into individual water baths for 60 min. Water samples were processed as described above. After completion of the water baths, frogs were immediately euthanized with an overdose of 20% benzocaine gel and rapidly decapitated. Trunk blood was collected into 1.5 mL Eppendorf tubes and centrifuged at 6000 rpm for 5 min (6-position rotor) to separate the plasma. Plasma volume was recorded, and samples were transferred into 1.5 mL eppendorf tubes prefilled with 750 uL of 96% ethanol. In the laboratory, testosterone was extracted from ethanol samples three times with freeze-decanting following the methodology in Goymann et al., (2007). Briefly, samples were dried down with N2 at 37°C. Dried pellets were resuspended in 4 mL of dichloromethane and 100 uL of distilled water and, then incubated at 4°C overnight for equilibration. The following day, samples were shaken for 1h and then centrifuged at 4000rpm for 10 min to separate the aqueous and organic phase, which was transferred into a new tube by freeze-decanting. This process was repeated twice, and the organic phase was then dried down at 37°C under N2 stream and then resuspended in the assay buffer supplied by the ELISA manufacturer and incubated overnight at 4°C.

### 2.3. Simulated territorial intrusion-tests (STIs)

#### 2.3.1. Field site and playback stimuli

Between February and April of 2018 and 2019, seventeen free-living adult *A. femoralis* males (mean ± SD = 2.95 ± 0.06 cm SUL) from a population located in the field station “Pararé” at Les Nouragues nature reserve in French Guiana (4°02’ N – 52°41’ W; Bongers et al., 2013) were used for the STI tests. STI tests consisted in presenting the playback of an artificial advertisement call featuring the spectral and temporal parameters of a nearby population within the nature reserve Les Nouragues (Gasser et al., 2009; Narins et al., 2003). To avoid pseudoreplication, we created 11 different playback stimuli (16-bit, 44.1-kHz WAV-file), which varied in the inter-note interval and the inter-call interval. Playbacks were broadcast using a loudspeaker (Creative MUVO 2c, Creative, Singapore) connected to a music player. Sound-pressure levels (SPLs) of every playback stimulus were calibrated at 75 – 80 dB using an SPL-meter (Voltcraft 329) at 1 m distance by adjusting the volume setting of the music player. All playbacks were conducted under rainless conditions and mostly between 14:00 and 17:00h.

#### 2.3.2. Experimental design

Focal males were tested using a pre-post experimental design which consists in comparing a hormonal and behavioural baseline with a post-social stimulation phase. During the baseline phase (A) we observed every focal male for 1 h from about 1.5 – 2 m distance and recorded the following behaviours: (1) duration of advertisement calls in seconds, (2) duration of “warm-up” calls in seconds (suboptimal advertisement calls of less than steady-state SPL; Jameson, 1954; Toledo et al., 2014), (3) duration of courtship calls in seconds, (4) # of feeding events, (5) # of head-body orientations (HBO) and (6) # of jumps. Observations were made between 08:00 and 18:00 h. We repeated the behavioural observations at least three times, at different times of the day (i.e. morning and/or afternoon), in non-consecutive days and with a minimum of three days in between observations. After every behavioural observation, each frog was gently captured with a plastic bag and immediately transferred into a water bath for 60 min to assess the baseline water-borne androgen concentration (A). Additionally, 24 females from the same population were also placed into individual water baths for 60 min in order to compare water-borne androgen baselines between sexes.

In the post-social stimulation phase (B), we confronted focal males to STIs exclusively when they were found calling. Once a focal male was located, we placed the loudspeaker on the forest ground at 1 – 1.5 m distance from the focal male. We considered a positive response (responding) when males approached the playback and reached a plastic-circular perimeter around the loudspeaker of 30 cm diameter (Amézquita et al., 2005). A negative response was recorded when males did not approach the loudspeaker and/or did not cross the perimeter before the playback was finished (non-responding). In order to determine the behavioural responsiveness of the frogs to the territorial challenge, we performed 3 STIs trials which were audio recorded and we measured the following behavioural parameters during each trial: (1) latency to the first head-body orientation towards the speaker, (2) latency to the first jump and, (3) latency until the frog reached the perimeter. Frogs were not handled or manipulated at least three days before any further STI. After the STIs (regardless whether the males responded or not) males were caught and immediately transferred to a series of three consecutive water baths of 60 min/each (1h, 2h and 3h). This sequence of water baths allowed us to determine a 3h timeline of androgen secretion in water. Time elapsed between the end of the STI and the beginning of the water baths was always less than 10 min. Water samples were collected individually after every 60 min water bath without manipulating the frog to avoid stress. For this, we used two flexible polymer tubing with one end attached to the glass box and the other end attached to a 20mL syringe. One tubing was used to pump the water into the glass box and the other was used to suck out the sample after every 1h water-bath. Samples were processed and extracted as explained in the water-borne extraction section. We repeated STIs three times per focal male with at least three days between trials.

### 2.4. Hormone assays

In order to estimate androgen concentration, we used a commercial enzymatic immunoassay for testosterone (ADI-900-065; Enzo Life Sciences, Farmingdale, NY, USA). Reconstituted samples were brought to room temperature and shaken at 500 rpm for 1 h prior the assay. Samples were plated in duplicate and assays were performed following the manufacturer’s protocol. Plates were read at 405 nm, with correction between 570 and 590 nm, using a microplate reader (Multiskan Go, Thermo Fisher Scientific Oy, Finland) and androgen concentrations were calculated using the Thermo Scientific SkanIt Software (version 4.1). The detection limit for the assay was 5.67 pg mL-1. The cross reactivity of the testosterone antibody with other androgens was below 15% (see manufacturers manual). The average intra- and inter-assay coefficient of variation were 3.38% and 11.05%, respectively.

### 2.5. Statistical analysis

Prior to analysis, hormone data were log-transformed to fit normality when necessary. In order to know whether water-borne androgen concentration was dependent on the frogs’ body size and/or body area, we first calculated the body area-SUL ratio for every frog. Later, we performed separate linear mixed models (LMM) for afternoon and morning baselines as response variables, body area-SUL ratio as fixed factor and the ID of the frogs as random factor. Because androgen baseline concentrations in the afternoon or the morning were not dependent on body area-SUL ratio (LMM: *β_morning_*= 1.18, *t*= 1.22, *P*= 0.22; *β_afternoon_*=1.68, *t*= 1.61, *P*= 0.11), water-borne androgen levels were not corrected for body size or area. In order to determine the release rate of androgens in water, we compared the time series water baths (15, 30 and 60 min) by performing a LMM to account for the repeated measurements, using the “lmer” function within the *lme4* package (Bates et al., 2015) in R (R Core Team, 2017). We used androgen concentration as the response variable, the time series of water baths as the fixed factor and the frog ID as the random factor.

To determine the parallelism between hormone concentration in blood and holding water, we performed a parametric correlation between the plasma and water-borne androgen concentrations using the Pearson’s product moment correlation coefficient. In order to compare water-borne androgen levels between males and females, we performed a two-sample t-test. Since time of the day, vocal and locomotor activity might be interdependent with androgen concentrations (Wada, 1986), we asked whether baseline androgen levels were dependent on natural behaviours and varied across the day. For this, we first performed a LMM with water-borne androgen levels as response variable, time of the day (morning/afternoon) as fixed factor and frog ID as the random factor. Then, we performed a Varimax normalized principal component analysis (PCA) in order to minimize redundancy among the behavioural variables by using the function “principal” within the R package *psych* (Revelle, 2019). Further, we performed a series of independent LMMs with the scores of the principal components obtained as response variables, time of the day (morning/afternoon) and water-borne androgen levels as fixed factors and frog ID as the random factor.

In order to investigate whether *A. femoralis* males respond to territorial challenges (STIs) with an increase in androgen levels, we first performed a LMM with androgen concentration as dependent variable, and the sampling time points (0h-morning/afternoon baselines-, 1h, 2h and 3h water bath sampling after STIs) as fixed effects. We used frog ID as the random factor to account for repeated measurements. In order to compare the androgen responsiveness to STIs between responding and non-responding males, we estimated the androgen responsiveness to male-male interactions (R_male-male_; Goymann et al., 2007) by computing the within-subjects standardized effect size (Cohen’s d) of the ratio between the water-borne testosterone concentration of every male after the STI and the baseline levels. Cohen’s d allows us to directly compare the magnitude of the androgen response by estimating the difference between pre (baseline) and post (STI-challenged frogs) water-borne androgen concentrations on a standardized scale (Goymann et al., 2007). For this, we used the function “cohens.d” within the R package *misty* (Yanagida, 2020).

Finally, in order to know whether the phonotactic approach of *A. femoralis* males is proportional with the androgen responsiveness, we first minimized redundancy among the three responsiveness latencies (latency to the first head-body orientation towards the speaker, latency to the first jump and, latency until the frog reached the perimeter) by using a varimax normalized principal component analysis (PCA). Then, we performed a LMM with the principal component scores as the response variable, the androgen responsiveness (R_male-male_) as the fixed effect predictor and the male ID as the random effect.

### 2.6. Ethics approval

All experiments were conducted in strict accordance with current Austrian, French and European Union laws and were approved by the Animal Ethics and Experimentation Board of the University of Vienna (No. 2018-010; 2019-002). Our study was approved by the technical director of the “Nouragues Ecological Research Station” where field work was conducted. We adhere to the “Guidelines for the use of live amphibians and reptiles in field and laboratory research” by the Herpetological Animal Care and Use Committee (HACC) of the American Society of Ichthyologists and Herpetologists. Collection permits were provided by the *Ministère de la transition écologique et solidaire*, *République Française* (No. TREL1902817S/152).

## 3. Results

### 3.1. Validation and sex differences of water-borne androgens

Recoveries of low and high standards were 98.21% and 105.98%, respectively. “Blank” water samples were below the detection limit of the assay (**Figure 1A**). Correlation between expected and obtained androgen concentrations in 2 mL aliquots was highly significant (*r*=0.99, *P*=0.006; **Figure 1B**), and falls within the range of detectability of the assay. Androgens released in 60 min water baths was significantly higher than 15 min (LMM: *β*=-112.43, *t*=-2.33, *P*=0.03; **Figure 1C**), but not than 30 min water baths (LMM: *β*=-46.05, *t*=-0.95, *P*=0.35; **Figure 1C**). Males had higher water-borne androgen levels than females in 60-min water baths (water-borne androgen mean ± SD: males=317.30±78.67 pg/mL; females=243.64±170.70 pg/mL; two sample t-test: *t*(33.51)= −3.07, *P*=0.004; **Figure 2**). Water-borne androgen concentration was positively correlated with plasma testosterone concentration (t=4.82, r=0.76, P<0.001; **Figure 3**).

**Figure 1.**
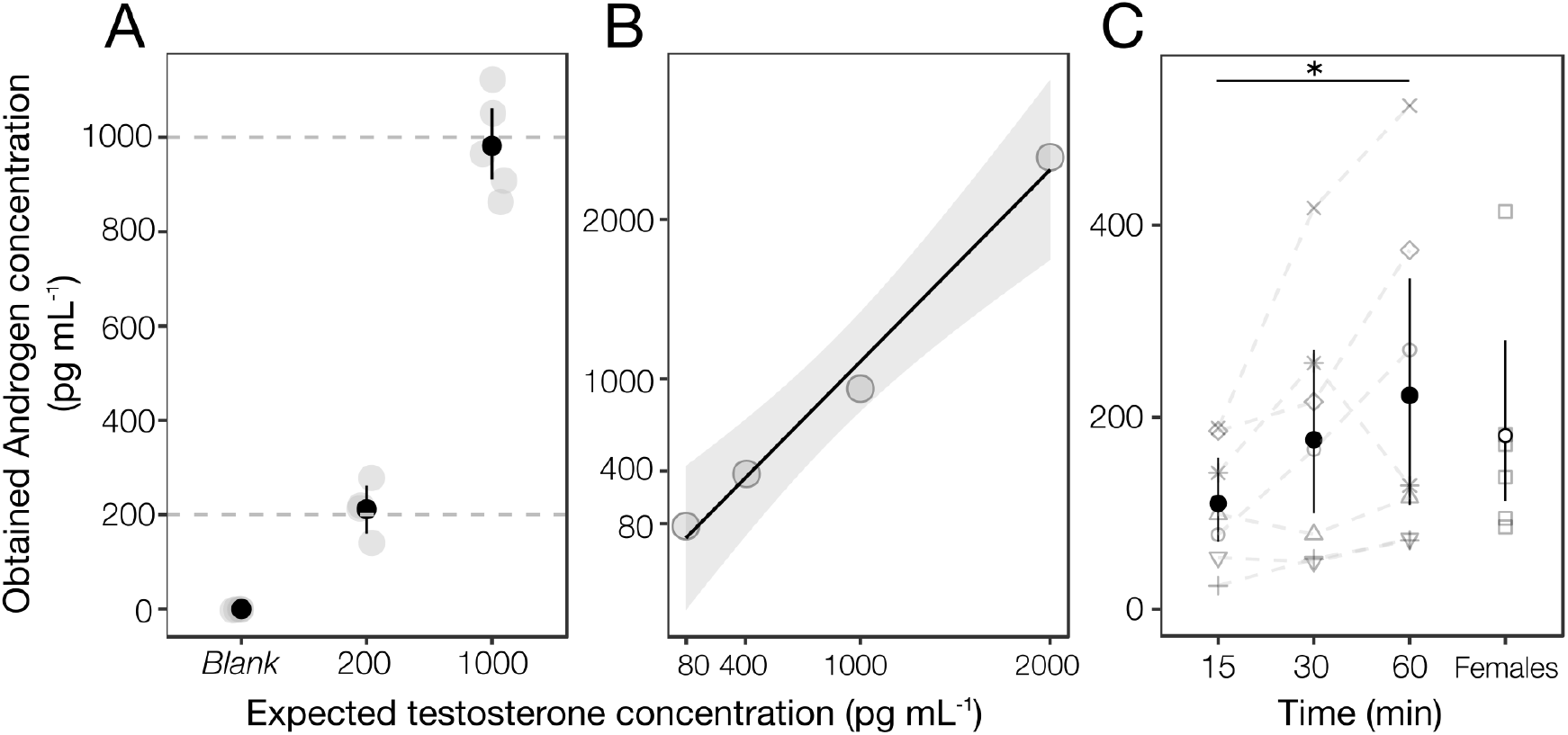
Validation of the extraction method for water-borne androgen. **(A)** Recovery percentages of testosterone standards of 0 (Blank), 200 and 1000 pg mL-1; **(B)** Correlation between expected and obtained testosterone concentrations in 2 mL aliquots; **(C)** Release rates of water-borne androgen in 60-min, 30-min and 15-min. Release rates are also shown for females in 60-min water baths. Solid black-points and bars represent the mean and 95% confidence intervals, respectively. *P<0.05

**Figure 2.**
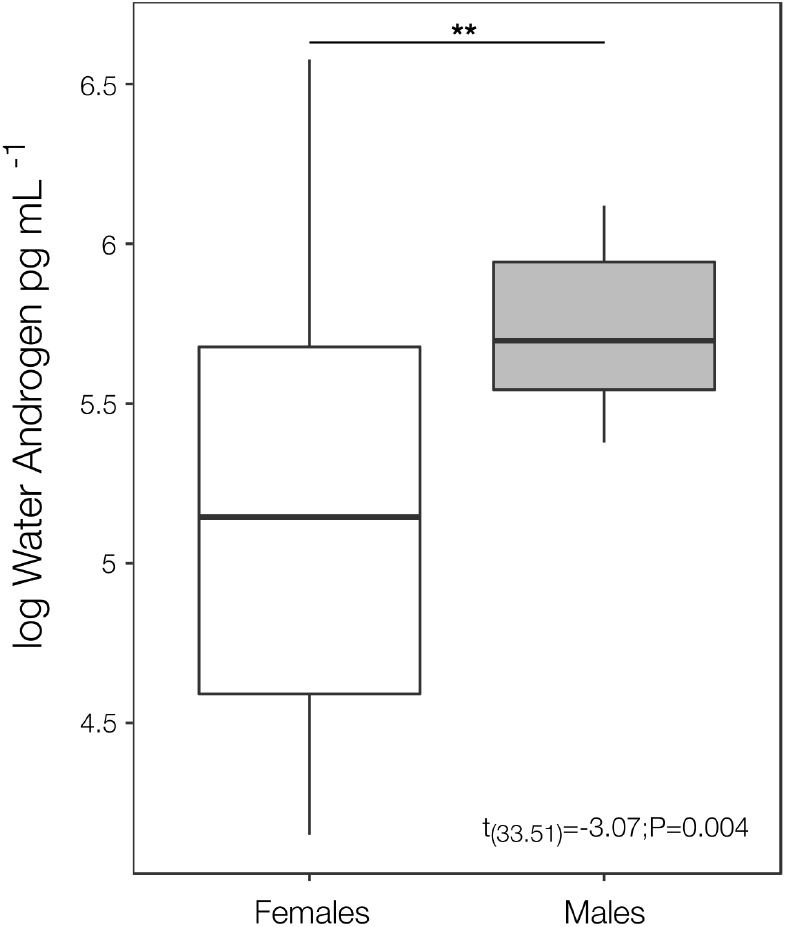
Comparison between males and females in water-borne androgen concentration. Both, males and females were placed in individual water baths for 60 min. **P<0.01

**Figure 3.**
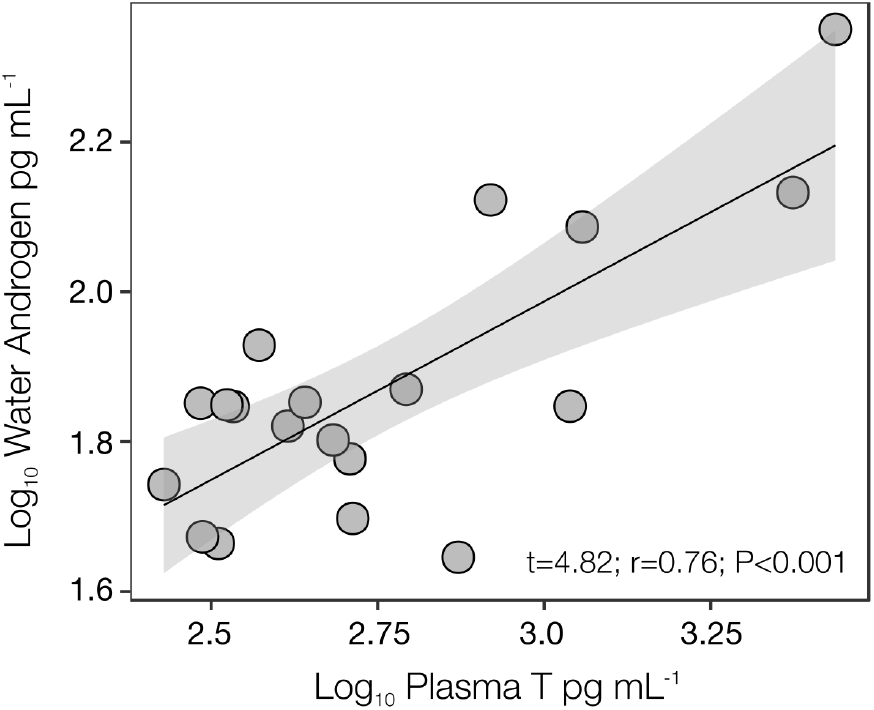
Parametric correlation (Pearson) between plasma testosterone and water-borne androgen concentrations. Shaded grey region represents 95% confidence intervals.

### 3.2. Daily variation of behaviours and water-borne androgen levels

Three components were generated with eigenvalues greater than 1 (**Table 1**): the first component (PC1) held 39% of the explained variance and represented positively vocal behaviour variables (advertisement and warm-up call durations). The second component (PC2) accounted for 31% of the source of variation and represented positively courtship behaviour variables (number of HBOs, jumps and duration of courtship calls). The third component (PC3) explained 30% of the variance and represented positively variables related to foraging behaviour (number of HBOs and feeding events).

**Table 1.**
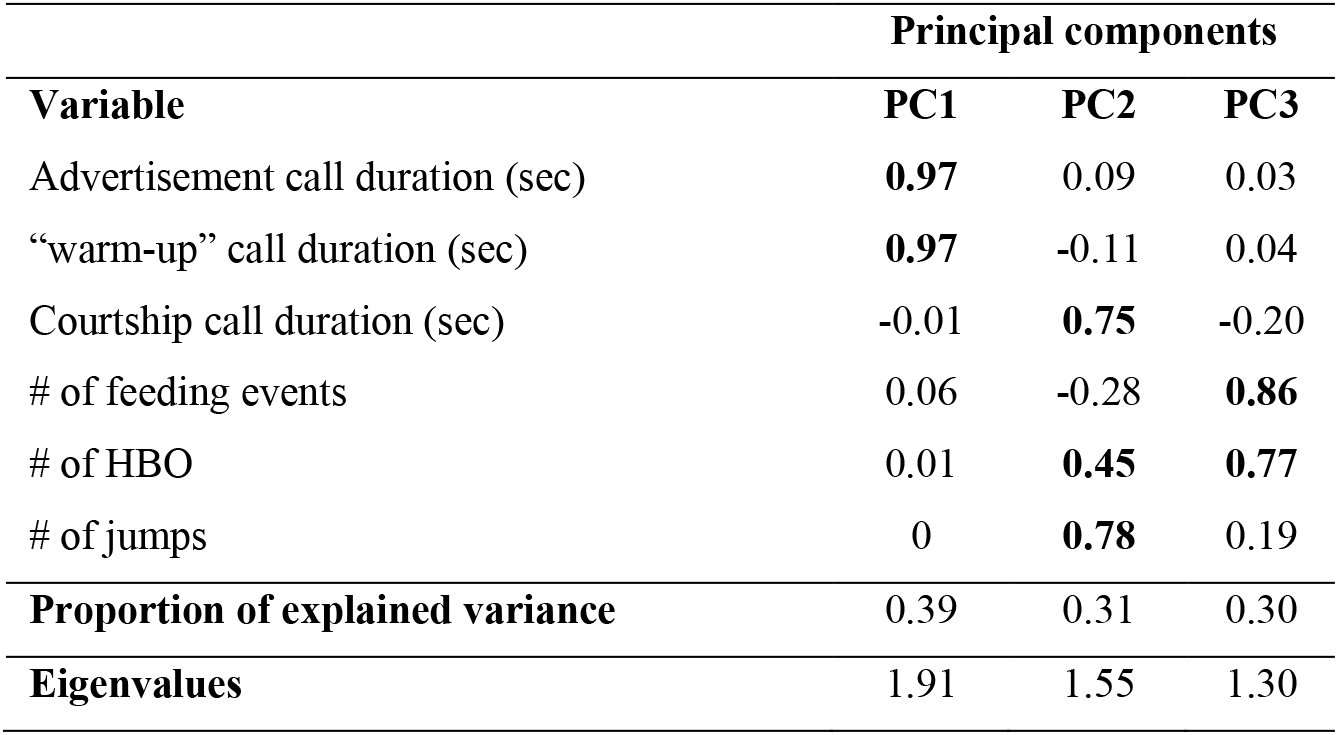
Principal Component Analysis showing the loadings matrix of the behavioural variables in principal components with eigen values greater than 1.

Baseline water-borne androgen concentrations and vocal behaviour (PC1) were significantly higher in the afternoon than in the morning (water-borne androgens: β=0.31, t=2.55, *P*=0.01; **Figure 4A;** PC1: *β*=0.56, t=3.62, *P*<0.001; **Figure 4B**). However, vocal behaviour was not dependent on water-borne androgen concentrations (PC1: *β*=0.07, t=0.42, *P*=0.67; **Figure 4B**). Courtship behaviour (PC2) and foraging behaviour (PC3) were not significantly different over the day (PC2: β=0.28, t=1.52, *P*=0.13; **Figure 4C;**PC3: *β*=-0.02, t=-0.1, *P*=0.92; **Figure 4D**) and/or dependent on water-borne androgen concentrations (PC2: *β*=0.1, t=0.55, *P*=0.58; **Figure 4C;** PC3: *β*=0.19, t=1, *P*=0.32; **Figure 4D**).

**Figure 4.**
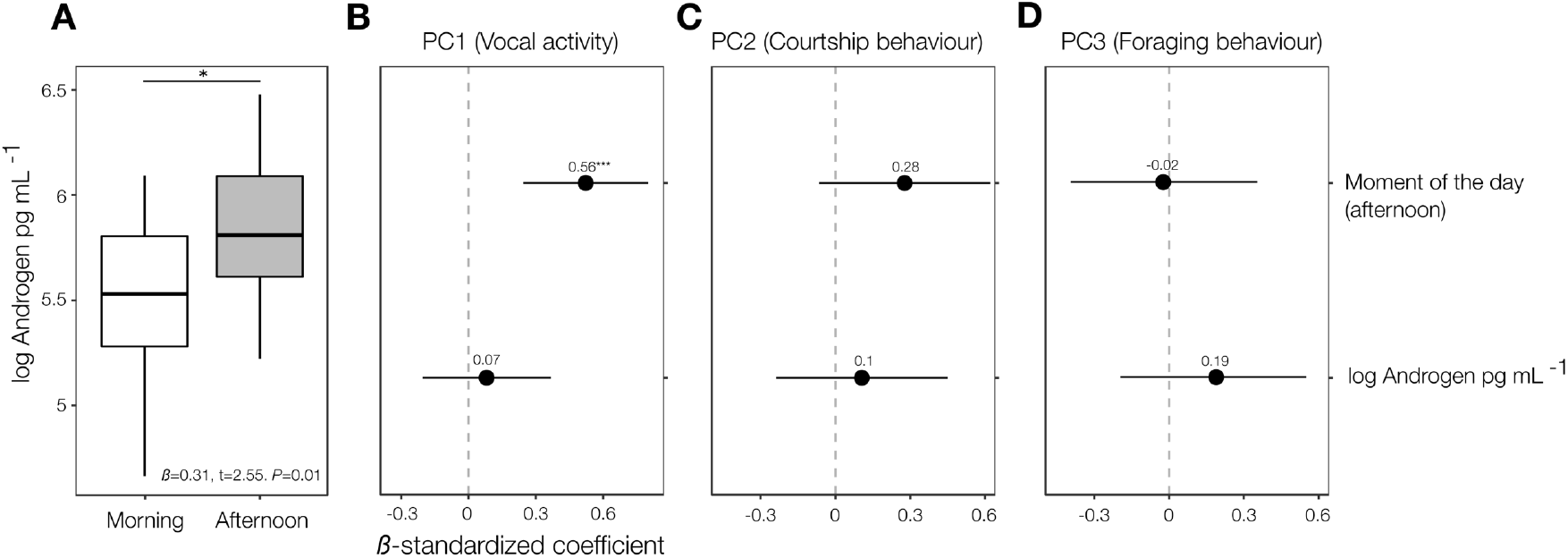
Water-borne androgen levels, vocal, courtship and foraging behaviour across the day. (A) Boxplot showing the difference of water-borne androgen concentration between morning and afternoon; Linear Mixed Model plots showing z-scores values (x-axis) and the effect size (numbers over mean-points) of time of the day and androgens over vocal activity (B), courtship behaviour (C) and foraging behaviour (D). Solid lines represent 95% confidence intervals. *P<0.05, **P<0.001.

### 3.3. Effect of STI on water-borne androgen levels (Rmale-male)

When frogs responded approaching towards the playback (i.e. positive phonotaxis), water-borne androgen levels significantly increased 1h after the STI compared to the baseline water-borne androgen levels in the morning (LMM: *β*=0.40, *t*=3.49, *P*=0.001; **Figure 5A**) and in the afternoon (LMM: *β*=0.21, *t*=0.10, *P*=0.04; **Figure 5A**). Subsequently, androgen concentration dropped nearly to the morning baseline in the 2h sampling point and under both, morning and afternoon baselines in the 3h sampling point (**Figure 5A**).

**Figure 5.**
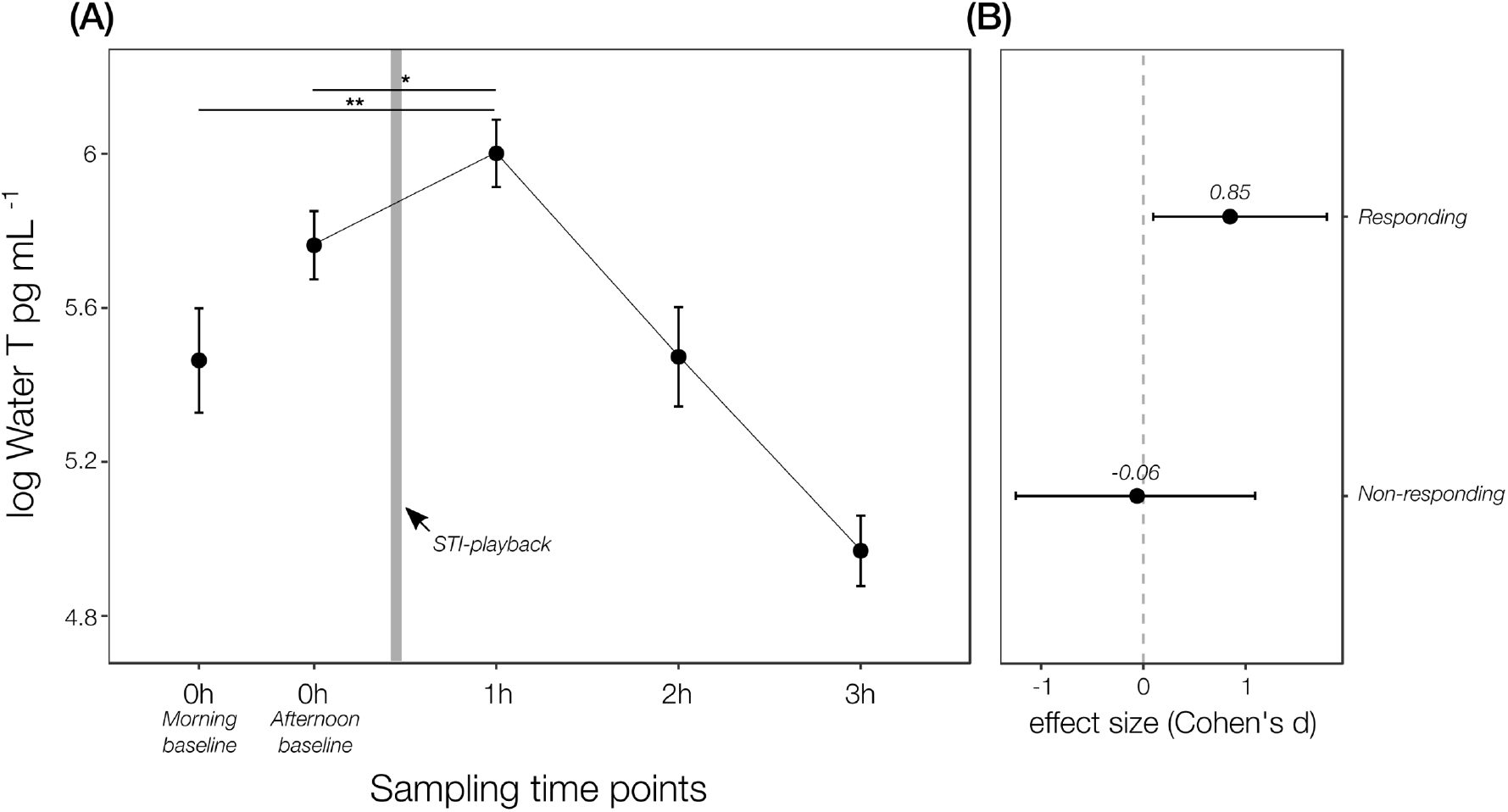
Androgen responsiveness after STI in *A. femoralis* males. **(A)** Comparison of baselines of water-borne androgen concentration (morning/afternoon; pre-STI) between concentrations over three sampling times (1h, 2h and 3h; post-STI). *P<0.05, **P<0.001. **(B)** Differences in effect size (±95% confidence intervals for both variables) of the male-male androgen responsiveness (dRmale-male) between responding (N=16) and non-responding (N=6) males.

Responsive males to the playback had a positive effect size and 95% confidence intervals crossed zero (Cohen’s d = 0.85 ± 0.94; **Figure 5B**), suggesting a positive effect of STI tests on water-borne androgen levels. On the other hand, non-responsive males had a negative (and close to zero) effect size and 95% confidence intervals did not cross zero (Cohen’s d = −0.06 ± 1.15; **Figure 5B**), and thus suggesting a null effect of STIs on androgen levels.

### 3.4. Effect of STI on the phonotactic behaviour

Three principal components were generated, but just one component with an eigen value greater than 1, which explained the 72% of the total variance (**Table 2**). This component represented positively the three responsiveness latencies (latency to the first head-body orientation towards the speaker, latency to the first jump and, latency until the frog reached the perimeter). The phonotactic approach of *A. femoralis* males towards the playback was not related to the androgen responsiveness (LMM: *β*=0.58, *t*=0.55, *P*=0.58).

**Table 2.** Principal Component Analysis showing the loadings matrix of three variables related to responsiveness latencies.

| Variable | Principal components |  |  |
| --- | --- | --- | --- |
|  | PC1 | PC2 | PC3 |
| Latency to the 1<sup>st</sup> head-body orientation (sec) | **0.99** | -0.01 | -0.16 |
| Latency to the 1<sup>st</sup> jump (sec) | **0.99** | 0 | 0.17 |
| Latency to the perimeter (sec) | **0.48** | **0.88** | 0 |
| **Proportion of explained variance** | **0.72** | 0.26 | 0.02 |
| **Eigenvalues** | **2.28** | 0.66 | 0.05 |

## 4. Discussion

Thirty years ago, an explanation for the facultative increase of males’ androgen levels in response to social challenges was named as the “Challenge Hypothesis” (Wingfield et al., 1990). Since then, numerous studies have been testing this hypothesis across different animal taxa with diverse life history. In this study we tested the Challenge Hypothesis in the highly territorial poison frog, *Allobates femoralis*. To do so, we compared males’ androgen concentrations quantified both in a non-stimulated condition (baseline) and following a simulated territorial intrusion (post-STI). We took advantage of water-borne hormones sampling, a non-invasive technique, to characterize androgen levels and could show that it closely reflects circulating plasma testosterone levels. Our results demonstrate that water-borne androgen increases after a STI in *A. femoralis* males only when males approached the playback loudspeaker. Therefore, our results provide novel support to the Challenge Hypothesis in a territorial frog. The intensity of the phonotaxis to the playback, however, was not related to males’ androgen responsiveness to STIs.

Water-borne androgen concentration significantly increased 1h after confronting *A. femoralis* males to STIs (i.e. playbacks). The Challenge Hypothesis predicts a short-term but distinct increase of androgen levels in response to social challenges (e.g. male-male competition). To the best of our knowledge, the only previous study that provides support for the challenge hypothesis in an amphibian species showed that males of the túngara frog (*Engystomops pustulosus*) increased water-borne testosterone after being challenged with a combined chemical (holding water containing excretions of conspecific calling males) and acoustic stimulus (Still et al., 2019). The general idea of the functional significance of the increase of testosterone above the breeding baseline levels is likely to prepare males for a potential agonistic encounter, such as by increasing its muscular contractile capacity and locomotor performance (Miles et al., 2007). In territorial species, this physiological boost is advantageous as it increases the chances of winning physical contests against intruders when competing for space and resources. Thus, androgen responsiveness to STIs in *A. femoralis* males is similar to that found in other vertebrates for which the Challenge Hypothesis is supported (reviewed by Moore et al., 2019).

As predicted by the Challenge Hypothesis, we observed a significant positive effect of STIs on water-borne androgen levels only in *A. femoralis* males that approached the playback, while those which did not approach also did not show an increase in water-borne androgen levels. Previous research in *A. femoralis* has proposed that males’ decision to approach an intruder and engage in a contest depends on whether the intruder represents a perceptible risk or not (Ursprung et al., 2009). The increased androgen levels might be consequent to the perception of an aggressive territorial intrusion, which might increase the likelihood to perform aggressive displays to repel the rival (Wingfield, 2005). However, other factors like the presence of another male (or a robotic decoy; Narins et al., 2003) or the motivational state of the challenged male might trigger a positive phonotaxis. Likewise, steroid hormones have been shown to act on brain areas related with the expression of the motivational state to approach and recognize competing signals (Oliveira, 2004; Adkins-Regan, 2005; Yao et al., 2008; Leary, 2009). Further research is needed to determine which factors influence the motivation to approach and attack an intruder in *A. femoralis*.

Despite a short-term increase in testosterone levels associated to a positive phonotaxis towards the loudspeaker, we did not find a relationship between androgen responsiveness and the latency of approach. In other words, males with higher androgen levels did not approach the STI faster. This may depend on the experimental setup of the STI and the nature of the STI stimuli (i.e. duration of the playback, live vs. synthetic decoy; reviewed by Goymann et al., 2007). For instance, an androgen response was only elicited in the túngara frog when chemical and acoustic stimuli were presented in combination (Still et al., 2019). Although *A. femoralis* males are strongly territorial and usually males jump towards the sound source in playback experiments (Hödl, 1983), they require to be confronted by bimodal signals (acoustic and visual) in order to display physical attacks (Narins et al., 2003). Thus, in *A. femoralis* males, playbacks alone may be enough to provoke a phonotactic reaction followed by an androgen response, but the intensity of phonotactic approach may depend on the combination of acoustic and visual signals (see also Sonnleitner et al. 2020). Further experiments on the hormonal and behavioural response to territorial intrusions in territorial frogs may profit from the combination of playbacks and robotic frog models in order to create more realistic situations.

We found higher water-borne androgen concentrations in males compared to females of *A. femoralis*. Androgens are the main class of sex hormones in male vertebrates and circulating androgens are typically lower in female vertebrates. In amphibians, several studies have found sex differences in plasma androgen levels, where males have higher circulating baseline concentrations than females. High androgen concentrations in male amphibians play a key role in the performance of territorial and reproductive behaviours such as vocal and clasping behaviours (Reviewed by Moore et al., 2005). However, it is noteworthy that hormonal differences between sexes are dynamic and change across behavioural contexts and at different life history stages (e.g. parental care, mating systems, sex-specific behaviours), rather than simply physiological differences settled through ontogeny (Adkins-Regan, 2005; Fischer and O’Connell, 2020). For instance, previous studies in other amphibian species have shown that females can show higher levels of androgens than males in relation to secretion of oestrogen, given that androgens are an obligate intermediate of oestrogen synthesis (i.e. aromatization; Delrio et al., 1979; Licht et al., 1983). In our study, we observed a large variation in water-borne androgen concentrations in *A. femoralis* females, with some individuals reaching even higher levels than males. At present, we can only speculate that such variation may be related to the breeding pattern of *A. femoralis*, which is an opportunistic breeder and males may have relatively low androgen concentrations throughout the year and not differ significantly from females outside reproductive periods.

The positive correlation between plasma and water-borne androgens in *A. femorali*s is in line with that found in other species e.g. fishes and amphibians (Baugh et al., 2018; Gabor et al., 2016, 2013; Kidd et al., 2010; but see Millikin et al., 2019 for non-correlation between water-borne and plasma corticosterone in spotted salamanders). Water-borne sampling has enormous advantages for estimating hormonal concentrations with little manipulation of the research animals (Narayan, 2013). Another benefit of water-borne sampling is that it can be performed repeatedly without harming the animal. For instance, researchers can evaluate hormone concentrations of individuals in different life history stages (Adkins-Regan, 2005; Baugh and Gray-Gaillard, 2020; Leary, 2009) and/or, compare hormonal responses between pre-and post-challenges (Bell, 2019; Still et al., 2019). Thus, in many cases water-borne sampling might even constitute a preferable alternative to invasive methods (i.e. blood sampling), offering new ways on how to study the interplay between social behaviour and hormones (Bell, 2019; Narayan, 2013; Wingfield et al., 2006).

We observed that water-borne androgen levels and vocal activity were higher in the afternoon than in the morning in *A. femoralis* males. In fact, previous research has found a higher calling activity peak of *A. femoralis* in the afternoon compared to the morning (between 1500-1730 h; Kaefer et al., 2012; Roithmair, 1992). *Allobates femoralis* males use advertisement calls to engage in social interactions with conspecifics (e.g. territory tenancy advertisement, inter-male spacing, courtship; Ringler et al., 2017; Rodríguez et al., 2020; Stückler et al., 2019). Interestingly, vocal behaviour was not dependent on androgen levels in *A. femoralis* males. Previous work reported positive association between testosterone and vocal behaviour in anuran amphibians (see below). Testosterone not only regulates the development of structures related to vocalization and neural pathways for the control of sound production (Reviewed by Leary, 2009; Moore et al., 2005), but also the motivation for calling and calling effort are androgen dependent in anurans (Burmeister and Wilczynski, 2001; Emerson and Hess, 1996; Solís and Penna, 1997). However, previous studies also showed that castrated and androgen treated males did not maintain or increase vocal behaviour (Burmeister and Wilczynski, 2001; Wetzel and Kelley, 1983), suggesting that androgens are needed but not the only hormones for eliciting vocal behaviour.

We found that elements related to courtship and foraging behaviour (e.g. # of head-body orientation, # of jumps, # of feeding events, courtship call duration) did not vary across the day in *A. femoralis* males. This result is not surprising, because courtship behaviour in *A. femoralis* males consists in a combination of acoustic cues (advertisement and courtship call) and a series of short locomotor events (courtship march), which usually start in the late afternoon (~17:15 h) and resume on the next morning (ending around 08:11 h; Stückler et al., 2019). Also, *A. femoralis* has a generalist feeding pattern and eats prey throughout the day (Pough and Taigen, 1990; Toft, 1980). Further, we found no relationship between courtship and foraging behaviour and water-borne androgen levels. Although vocal and courtship behaviours are androgen-dependent in most acoustically communicating species, the expression of socially evoked behaviours in other anurans depend also on other hormones such as neuropeptides, and/or the interaction between both classes of hormones (reviewed by Moore et al., 2005). For instance, the neuropeptide arginine vasotocin influences the motivation to call and courtship behaviour in frogs and salamanders (Burmeister and Wilczynski, 2001; Leary, 2009; Propper and Dixon, 1997), and at the same time its concentration in the brain depends on androgen concentrations (Boyd, 1994). The synergistic effects of androgen hormones and neuropeptides on courtship behaviour need further investigation in poison frogs.

Water-borne androgen was increased 1h after the STI but returned to baseline levels 2h after the STI. There are at least two possible reasons for such a pattern. First, short-term changes in androgen levels in non-seasonal breeders have been associated with the trade-off between parental care and aggressiveness (Wingfield et al., 1990). In other words, androgen levels can facultatively rise during male-male contests but decrease when males are parenting the broods. *Allobates femoralis* males typically perform tadpole transport and, although we were unable to evaluate the effect of parental care before or after presenting the STIs, unpublished data suggest that parenting males have significantly lower water-borne androgen compared to non-parenting males (Rodríguez et al., unpublished data). Second, there are costs associated with maintaining high androgen levels for a prolonged period of time such as the suppression of immune function, increasing of parasitic infections (Folstad and Karter, 1992) and impairing parental care (Wingfield et al., 1990). Thus, the return of androgens to baseline levels after a short-term increase may ease the resume of ongoing activities just before the intrusion. Interestingly, water-borne androgen levels went below the pre-STI baseline levels 3h after the STI. This reduction might be the consequence of negative feedback of the hypothalamo-pituitary-interrenal (HPI) axis (Yao & Denver, 2007). Additionally, we cannot exclude that there are some inhibitory effects caused by a stress response resulting from the isolation of the frogs for a prolonged period of time in the glass box. Additional research is necessary to further investigate these questions in *A. femoralis*.

## 5. Conclusions

Our study is one of the first to support the Challenge Hypothesis in a territorial frog, by using STIs and a non-invasive technique to characterize androgen levels. We found that water-borne androgen is responsive to social challenges in males of the highly territorial poison frog, *Allobates femoralis*. Since water-borne hormones provide biologically and physiologically relevant information by mirroring hormone levels in plasma, the integration of territorial intrusion experiments and non-invasive hormone sampling may allow researchers to test the “Challenge Hypothesis” in animal systems with a broad suite of life histories.

## Acknowledgements

We thank the Nouragues research field station (managed by CNRS) which benefits from “Investissement d'Avenir” grants managed by Agence Nationale de la Recherche (AnaEE FranceANR-11-INBS-0001; Labex CEBA ANR-10-LABX-25-01). This work was funded by the Austrian Science Fund (FWF): W1262-B29. We specially thank the Vienna Zoo (Schönbrunner Tiergarten Ges.m.b.H), the Austrian Herpetological Society (Österreichische Gesellschaft für Herpetologie) and the Centre National de la Recherche Scientifique (CNRS) for additional funding provided to CR. We are especially grateful with P. Gaucher and N. Marchand for the invaluable logistic support. For providing field assistance we are grateful with A. Lembens, K. Dellefont, C. Leeb and M. Peignier.

## Notes

### Competing Interest Statement

The authors have declared no competing interest.

